# Deep, unbiased and quantitative mass spectrometry-based plasma proteome analysis of individual responses to mRNA COVID-19 vaccine

**DOI:** 10.1101/2024.04.22.589104

**Authors:** Ting Huang, Alex Rosa Campos, Jian Wang, Alexey Stukalov, Ramón Díaz, Svetlana Maurya, Khatereh Motamedchaboki, Daniel Hornburg, Laura R. Saciloto-de-Oliveira, Camila Innocente-Alves, Yohana P. Calegari-Alves, Serafim Batzoglou, Walter O. Beys-da-Silva, Lucélia Santi

## Abstract

Global campaign against COVID-19 have vaccinated a significant portion of the world population in recent years. Combating the COVID-19 pandemic with mRNA vaccines played a pivotal role in the global immunization effort. However, individual responses to a vaccine are diverse and lead to varying vaccination efficacy. Despite significant progress, a complete understanding of the molecular mechanisms driving the individual immune response to the COVID-19 vaccine remains elusive. To address this gap, we combined a novel nanoparticle-based proteomic workflow with tandem mass tag (TMT) labeling, to quantitatively assess the proteomic changes in a cohort of 12 volunteers following two doses of the Pfizer-BioNTech mRNA COVID-19 vaccine. This optimized protocol seamlessly integrates comprehensive proteome analysis with enhanced throughput by leveraging the enrichment of low-abundant plasma proteins by engineered nanoparticles. Our data demonstrate the ability of this nanoparticle-based workflow to quantify over 3,000 proteins from 48 human plasma samples, providing the deepest view into COVID-19 vaccine-related plasma proteome study. We identified 69 proteins exhibiting a boosted response to the vaccine after the second dose. Additionally, 74 proteins were differentially regulated between seven volunteers, who contracted COVID-19 despite receiving two doses of the vaccine, and the ones who did not contract COVID-19. These findings offer valuable insights into individual variability in response to vaccination, demonstrating the potential of personalized medicine approaches in vaccine development.

## Introduction

Prompted by the recent global COVID-19 pandemic, nanomedicine-driven vaccines emerged as pivotal agents in preventive and therapeutic interventions^1^. However, clinical studies have shown that immune responses to these vaccines differ among individuals^2,3^. Factors such as genetics, health status, and environmental conditions contribute to this diversity in responses. These variations could often by tracked down to specific molecular patterns in each individual. Here we focus on the proteins found in blood plasma at each stage of vaccination. Plasma proteins such as immunoglobulins, cytokines, chemokines, and those involved in complement pathways, serve as indicators of immune status and reaction to the vaccine^4,5^. The degree of protection afforded by the vaccine is directly influenced by the concentration of these proteins; while some individuals gain full benefit, others do not respond as effectively^6^. Comprehensive analysis of plasma proteome thus offers valuable insight into how the body responds to vaccination and provides the gateway towards personalized vaccination strategies.

Plasma proteome is accessible and thus enables detailed longitudinal studies of health status and disease progression^7,8^. Particularly, plasma proteomics has yielded novel insights into vaccine responses and the molecular fingerprints of immunization^6,9,10^. Despite its biological and clinical utility, plasma proteomics trails behind genomics in widespread clinical and translational research primarily due to a more diverse biochemistry of proteins and technical challenges in detecting low abundant proteins. Recent advancements in measuring plasma proteome encompass two types of methods. Targeted, affinity-based methods facilitate the measurement of a pre-defined list of proteins^11^. Conversely, untargeted methods, mainly using liquid chromatography-tandem mass spectrometry (LC-MS/MS) analysis, aim to provide a quantitative snapshot of total proteome^12^. However, traditional LC-MS/MS methods face limitations in either proteome coverage or throughput due to the vast dynamic range of protein concentrations in plasma, where high abundance proteins often suppress the signal of the low abundant ones.

Our study leverages a novel nanoparticle-based untargeted LC-MS/MS workflow, which has shown exceptional capacity for in-depth plasma protein coverage, quantitative performance, and analysis throughput^8,13–16^. Here, we applied this workflow to assess the proteomic changes in a cohort of 12 volunteers following two doses of the Pfizer-BioNTech mRNA COVID-19 vaccine. In total, we identified 3,094 proteins in this cohort – a threefold improvement in coverage compared to the previous COVID-19 inactivated vaccine study that employed a serum depletion workflow ^6^. Thereby our study provides the deepest plasma proteomic data on the mRNA COVID-19 vaccination to date. These data revealed 69 proteins differentially regulated in a dose-dependent way in response to vaccination, as well as 71 proteins that are specifically regulated in participants that contracted COVID-19 after vaccination, thereby emphasizing the importance of global proteome profiling in understanding the molecular basis of vaccine response and efficacy.

## Experimental Section

### Sample Collection

This study was conducted in accordance with the Declaration of Helsinki and approved by the Brazilian National Review Board (Comissão Nacional de Ética em Pesquisa, CAAE number 50100321.8.0000.5347). Samples were collected from 12 volunteers ranging from 20 to 30 years old, between July and October 2021 after signed consent. The group consisted of 4 male and 8 female participants with a 23.6 average age. All participants received two doses of COVID-19 mRNA Pfizer-BioNTech vaccine. Blood samples were collected at 4 timepoints: before vaccination (Sample PreVax1), within 24 hours after the first dose (Sample PostVax1), before the second dose (Sample PreVax2) and within 24 hours after the second dose (Sample PostVax2) (**Figure 1A** and **Supplementary Table 1**). The second dose was around 60 days after the first dose. A total of 48 plasma samples were collected and stored at –80°C until use. Among 12 participants, 2 participants (S1 and S11) had a history of COVID-19 before or during sample collection, while others did not.

Several months after vaccination, follow-up surveys were carried out by July 2022 to inquire whether the volunteers had tested positive for COVID-19 and, if so, to document their symptoms. Based on the survey results, the 12 volunteers were categorized into two subgroups: (I) NONCOVID (n=5), comprising those who either did not test positive or did not undergo a COVID test, and (II) COVID (n=7), consisting of individuals who tested positive after completing the two-dose vaccination. One NONCOVID individual (S7) reported symptoms but did not undergo a COVID test for confirmation.

### Automated Plasma Sample Processing with Proteograph^TM^ Workflow

Plasma samples were processed with the SP100 automation instrument and the Proteograph™ Assay Kit (Seer, Inc.), employing five distinctly functionalized nanoparticles (NPs) within a fully automated workflow (**Figure 1A**). A total of 250 µL of plasma was equally aliquoted into five tubes, each containing 40 µL of plasma subjected to one-hour incubation with the proprietary NPs provided in the Proteograph Assay Kit (v1.2). The incubation step facilitated protein corona formation on NP surfaces and was followed by a series of gentle washes to eliminate non-specific and weakly bound proteins. Subsequently, proteins bound to the NPs underwent reduction, alkylation, and digestion with Trypsin/Lys-C, yielding tryptic peptides for downstream LC-MS/MS analysis. The peptides were then desalted, and all detergents were removed using a mixed media filter plate and a positive pressure (MPE) system. The resulting peptides were eluted into a deep-well collection plate in a high-organic buffer. Immediately after peptide elution, a peptide quantitation assay was conducted using the Pierce Fluorescent Assay Kit to determine the peptide yield for each well.

### Sample Multiplexing with TMTpro 18-plex Labeling

The tryptic peptides derived from the five nanoparticles were combined into a single sample for subsequent TMT labeling (**Figure 1A**). Each pooled peptide sample was dried in a SpeedVac for 3 hours and then reconstituted directly in a solution consisting of 50% acetonitrile in 100mM HEPES (pH 8) containing the TMT tags from the TMTpro 18-plex reagent (Thermo Fisher Scientific). The peptide-TMT mixtures were incubated in a thermomixer for 1 hour at 25°C and 600 rpm, with the reaction being halted by the addition of 2% hydroxylamine to achieve a final concentration of 0.2%. This mixture was further incubated for 15 minutes at 25°C and 600 rpm. Labeled peptides for each 18-plex batch were pooled and dried using a SpeedVac system, followed by reconstitution in 0.1% formic acid (FA) for desalting using a C18 TopTip (PolyLC, Columbia, Maryland) according to the manufacturer’s guidelines. The 48 plasma samples obtained from 12 volunteers were then allocated into three TMT batches, ensuring that all four samples from the same individual were assigned to the same batch. Furthermore, two TMT batches (sets #1 and #2) were comprised by female-only samples, and one batch (set #3) was male-only samples.

Desalted TMT-labeled peptide pools were dried in a SpeedVac system before reconstitution in 20 mM ammonium formate (pH ∼10) for chromatographic fractionation. This chromatographic separation was accomplished using a Waters Acquity BEH C18 column (2.1x 15 cm, 1.7 µm pore size) mounted on an M-Class Ultra Performance Liquid Chromatography (UPLC) system (Waters). The peptide elution was carried out through a 35-minute gradient: 5% to 18% B in 3 min, 18% to 36% B in 20 min, 36% to 46% B in 2 min, 46% to 60% B in 5 min, and 60% to 70% B in 5 min (A=20 mM ammonium formate, pH 10; B = 100% ACN). A total of 48 fractions were collected and non-contiguously pooled into 24 fractions, which were then dried completely in a SpeedVac concentrator, prior to analysis.

The dried peptide fractions were reconstituted with 2% ACN and 0.1% FA, and subjected to Reversed Phase (RP) LC-MS/MS analysis using an EASY-nLC^TM^ 1200 system (Thermo Fisher Scientific) coupled to an Orbitrap LumosTribrid^TM^ mass spectrometer equipped with FAIMS Pro™ Interface (Thermo Fisher Scientific) utilizing a two-hour gradient. Peptides were separated using an analytical C18 Aurora^TM^ column (75µm x 250 mm, 1.6µm particles; IonOpticks) at a flow rate of 300 nL/min with an 80-minute gradient: 1% to 6% B in 0.5 min, 6% to 23% B in 50 min, 23% to 34% B in 29 min, and 34% to 48% B in 0.50 min (A= FA 0.1%; B=80% ACN: 0.1% FA).

The mass spectrometer operated in a positive data-dependent acquisition mode, employing the FAIMS Pro Interface device at standard resolution with the temperature of both FAIMS inner and outer electrodes set to 100°C. A “three MS experiments” method was configured, wherein each experiment employed distinct FAIMS compensation voltage: −45, −65, and −80 Volts, respectively, each with a 1-second cycle time. A high-resolution MS1 scan was conducted in the Orbitrap (m/z range 350 to 1,500, 60k resolution, AGC 4e5 with a maximum injection time of 50 ms, RF lens at 30%) in top-speed mode with a 1-second cycle time for the survey and the MS/MS scans. For MS/MS spectra, ions with a charge state between +2 and +7 were using a 0.7 m/z isolation window isolated with the quadrupole mass filter. Subsequently, they were fragmented with higher-energy collisional dissociation (HCD) using a normalized collision energy of 35%. The resulting fragments were detected in the Orbitrap at 50k resolution, AGC of 5×10⁴, and a maximum injection time of 86 ms. The dynamic exclusion was set to 20 sec with a 10 ppm mass tolerance around the precursor.

### Database Search for Proteomics Quantification

All mass spectra files were analyzed with SpectroMine^TM^ software (Biognosys, version 2.7.210226.47784) using the TMTpro^TM^ 18-plex default settings. The spectral data were searched against the UniProtKB/Swiss-Prot human protein sequence database (January 2022). The search criteria were set as follows: full tryptic specificity was required (cleavage after lysine or arginine residues unless followed by proline), up to two missed cleavages were allowed, carbamidomethylation (C), TMTpro (K and peptide N-terminus) were set as fixed modification and oxidation (M) as a variable modification. The false identification rate was set to 1% at PSM, peptide and protein levels. PSM report was exported from SpectroMine for further analysis (data available in ProteomeXchange XXX).

We utilized the R package MSstatsTMT (version 2.2.7) to conduct global median normalization on peptide intensity data, perform fraction aggregation, quantify proteins, and implement per-protein local normalization based on two pooling samples ^17,18^. The normalized protein abundance table is available in **Supplementary file 1**. We then used the web server DAVID^19^ to generate the list of Gene Ontology Biological Processes (GOBP) pathways enriched among the identified proteins in comparison to the total searched proteome.

### Differential Abundance and Pathway Enrichment Analysis

Differential protein abundance analysis was carried out using MSstatsTMT-based statistical test between all pairwise combinations of the four timepoints (PreVax1, PostVax1, PreVax2, and PostVax2), considering ALL individuals, NONCOVID individuals, and COVID individuals separately. The resulting p-values were adjusted for multiple testing by the Benjamini-Hochberg method. A protein was considered significantly differentially abundant if its adjusted p-value was below 0.05.

For each pair of the time points, we used the fgsea package^20^ to perform Gene Set Enrichment Analysis (GSEA)^21^ on proteins ranked by their fold-changes in our differential analysis, considering ALL individuals, NONCOVID individuals, and COVID individuals separately. We defined GO Biological Process term as significantly enriched, when its Benjamini-Hochberg-adjusted p-value was less than 0.05. The pathway enrichment analysis results are available in **Supplementary file 3**.

## Results

### Nanoparticle-based workflow with deep proteome coverage

In total, nanoparticle-based workflow combined with TMTpro 18-plex allowed us to quantify 23,372 peptides and 3,094 protein groups in 48 plasma samples from 12 individuals across 4 time points (**Figure 1A**). 3,068 of the 3,094 proteins had 10% or fewer missing values. Compared with the previous COVID-19 inactivated vaccine study^6^ that employed a serum depletion workflow, our nanoparticle-based approach expanded proteome coverage by threefold under the same 10% missingness criterion. We detected around 80% of the proteins reported by the previous study, as well as additional 2,305 unique proteins (**Figure 1B**). We used the Human Plasma Proteome Project (HPPP) database^22^ to estimate the absolute concentration of identified proteins and observed that our workflow dramatically enhances the detection of low-abundant proteins (**Figure 1C** and **Supplementary Figure 1**). The increased depth of proteome analysis allowed us to cover a diverse set of biological pathways. The top 15 Gene Ontology Biological Processes (GOBP), that are enriched among the identified proteins, spanned various immune-related pathways, which are crucial to understand vaccine response (**Figure 1D** and **Supplementary Figure 2**). Our study identified 229 cytokines and protein hormones, also improving the coverage of these important low-abundant immunity-related proteins in comparison to the previous studies (**Supplementary Figure 3**).

**Figure 1:**
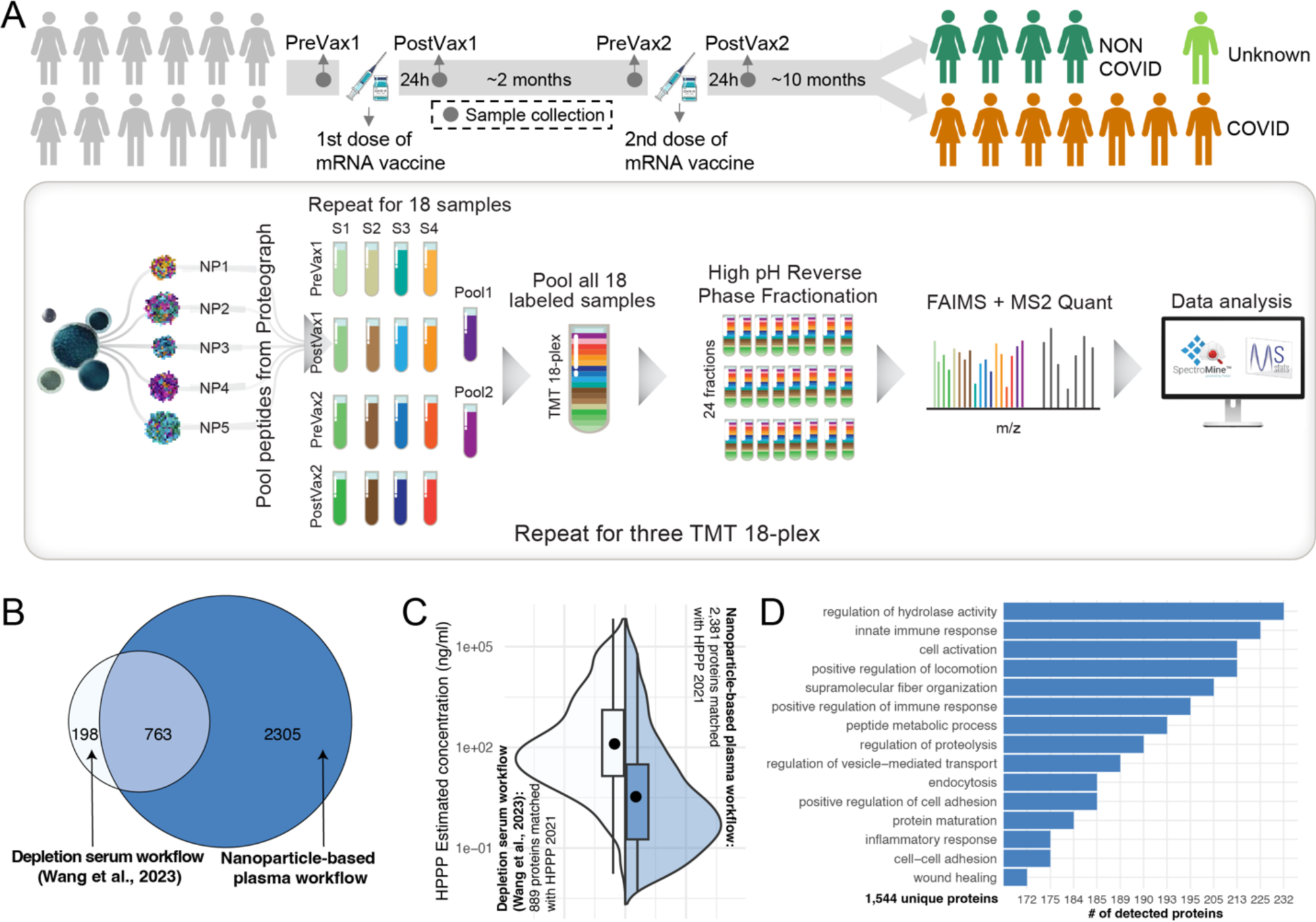
Overview and performance of nanoparticle-based workflow. (A) Experimental Workflow: 48 plasma samples were obtained from 12 volunteers before and after they received the first and second doses of the Pfizer-BioNTech mRNA COVID-19 vaccine. Follow-up surveys revealed that 7 volunteers, categorized as COVID, tested positive even after receiving both vaccine doses. Another 5 volunteers, labeled as NONCOVID, either tested negative throughout the study period or did not take a COVID test. One participant, classified as UNKNOWN, experienced symptoms suggestive of COVID-19 but did not confirm their status with a test. Plasma samples from all volunteers were processed using the SP100 automation instrument with the Proteograph Assay kit (Seer, Inc.), featuring five functionally distinct nanoparticles (NPs). Tryptic peptides from these NPs were pooled for each single sample for TMT labeling. The 54 pooled samples were divided into three TMTPro 18-plexes, each including PreVax1, PostVax1, PreVax2, and PostVax2 samples from four individuals, along with two pooled samples. Following TMT labeling and high pH reverse phase fractionation of peptides, samples were analyzed by LC-MS/MS using an Orbitrap Fusion Lumos MS equipped with FAIMS Pro Interface (Thermo Fisher Scientific). Raw MS spectra were processed using SpectroMine^TM^ (Biognosys), and the R/Bioconductor package MSstatsTMT was used to detect differentially abundant proteins. (B) Comparing the number of proteins identified by our nanoparticle-based plasma workflow and the existing COVID-19 inactivated vaccine study^6^ using a depletion serum workflow. Protein counts are reported at 10% missingness cutoff. (C) Comparing the sensitivity of low-abundant protein detection using estimated plasma protein concentrations from the Human Plasma Proteome Project (HPPP) database ^22^. (D) Top 15 GO Biological Processes enriched among the proteins identified in our study.

### Measuring plasma proteome changes after two-dose mRNA vaccination

To evaluate global proteomic responses to a two-dose mRNA COVID-19 vaccine, we conducted a differential analysis comparing pre- and post-vaccination protein levels across all participants (**Figure 2A**). A total of 69 proteins exhibited significant changes after the two vaccine doses (PreVax1 vs PostVax2), comprising 32 up-regulated and 37 down-regulated proteins. Interestingly, for most of the upregulated proteins, their response followed a “N” pattern: the initial quick increase of the abundance after the first dose (PostVax1 timepoint), followed by the decline towards the baseline after 60 days (PreVax2 timepoint), and then even more substantial increase after the second dose (PostVax2 timepoint). Conversely, most down-regulated proteins exhibited a continuous decrease in expression throughout the two doses of vaccine, with only five proteins following a reversed “N” pattern, suggesting that different mechanisms of regulation might be employed.

**Figure 2:**
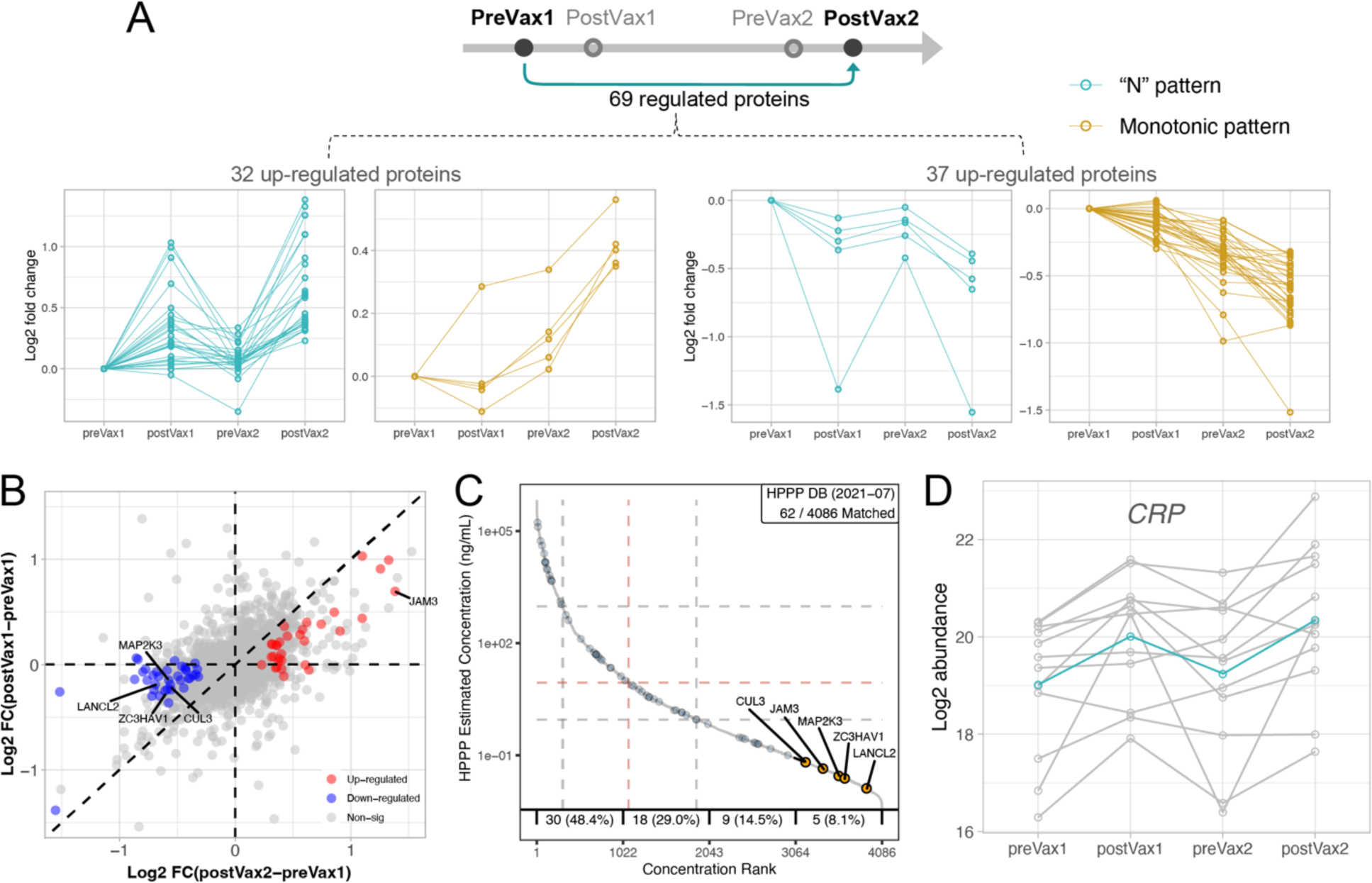
Analysis of protein regulation after the 2 doses of mRNA COVID-19 vaccine in all 12 Individuals. (A) Comparison of significantly regulated proteins (adjusted p-value ≤ 0.05) between PreVax1 and PostVax2 time points for all 12 individuals. The line plots show the dynamics of the 69 significantly regulated proteins relative to their PreVax1 levels. Most of upregulated proteins display an “N” pattern (leftmost; in teal), whereas those downregulated proteins typically show a consistent decline in expression throughout the two vaccine doses (rightmost; in amber). (B) Comparison of protein log₂ fold changes between PostVax2 and PreVax1 (X axis) and between PostVax1 and PreVax1 (Y axis). Red color highlights 32 significantly up-regulated proteins, and blue color highlights 37 significantly down-regulated proteins. The five lowest concentrated proteins among the 69 proteins are labeled. (C) Plasma protein concentrations (as reported by the HPPP database) of the 69 differentially regulated proteins. 14 proteins are from the lower concentration range (concentration rank between 2043 and 4086), with the 5 lowest concentrated proteins (concentration rank between 3064 and 4086) highlighted in yellow. (D) The dynamics of Serum C-reactive Protein (CRP) abundance. Each grey line represents an individual, and the teal line indicates the mean intensities across all 12 individuals.

While statistically significant changes in the abundance of 69 proteins were detected following the second vaccine dose, no statistically significant changes were observed after the first dose. However, for most of the proteins, the trend in abundance changes was consistent after both doses, though the extent of these changes was amplified following the second dose (**Figure 2B**). This observation suggests that these proteins are regulated already after the initial vaccination, but the changes do not reach statistical significance, likely due to the study size.

These differentially regulated proteins spanned the full dynamic range of plasma concentrations, as estimated by the Human Plasma Proteome Project database (HPPP)^22^ (**Figure 2C**). Notably, the five lowest concentrated plasma proteins (CUL3, JAM3, MAP2K3, ZC3HAV1, LANCL2) give important insights to understand the organism response to COVID-19 mRNA vaccine. The downregulation of MAP2K3, a kinase involved in activating p38 MAPK, which in turn regulates the production of pro-inflammatory cytokines in response to the vaccine^23^, may play a role in mitigating excessive inflammation and reducing symptoms severity. Similarly, modulation of LANCL2, known for its anti-inflammatory effects post-viral infection^24^, could modulate the immune response to vaccination. Interestingly, the upregulation of JAM3, recognized for its role in leukocyte trafficking and inflammation^26^, suggests a potential mechanism by which immune responses to the vaccine are enhanced through improved immune cell migration. Additionally, ZC3HAV1 is an antiviral protein activated by interferons that can limit SARS-CoV-2 replication^27,28^. Furthermore, the involvement of CUL3 in regulating T follicular helper (TFH) cell differentiation^29^, was recently identified in the context of SARS-CoV-2^30^, suggests a mechanism for long-term protective immunity, indicative of vaccination success. While our study was not designed to elucidate the specific molecular mechanisms of their regulation, the fact that we consistently observe the regulation of these low-abundant immunity- and inflammation-related proteins is intriguing and suggests further exploration.

Serum C-reactive protein (CRP) was the differentially expressed protein with the highest abundance (**Figure 2D**). This liver-produced protein, which serves as an early marker of infection and inflammation, was already identified as a robust biomarker of severe COVID-19 cases^31^. Interestingly, while the baseline abundance of CRP varied between participants, its dynamics consistently followed the same “N” pattern across participants.

### Plasma proteome profiling of NONCOVID participants

To further study the individual regulation of plasma proteome in response to vaccination, the cohort was stratified into COVID and NONCOVID groups. The COVID group comprised of participants who tested positive within ten months after completing the two-dose vaccination, the other participants were assigned to NONCOVID group. Among the 69 differentially abundant proteins, most have a larger magnitude of change in the NONCOVID group compared to COVID group (**Supplementary Figure 4**). This suggests that stronger regulation of these proteins in the NONCOVID group may contribute to a more robust immune protection.

When differential abundance analysis was performed separately within each group, 77 proteins showed significant changes across different time periods in NONCOVID group, while only 4 proteins showed significant changes in COVID group, and 76 in ALL participants (**Figure 3A**). Only 16 of 77 proteins differentially regulated in NONCVOID group were among the proteins regulated in ALL participants (see **Supplementary Figure 5**). We observed that 66 of 77 proteins (86%) were downregulated in NONCOVID group, while no such bias was observed in ALL participants with only 39 of 76 proteins (51%) downregulated. Furthermore, within ALL group, only 3 proteins displayed statistically significant regulation before the second dose was administered (PreVax2 vs PreVax1 comparison), whereas 62 proteins exhibited significant regulation in NONCOVID group between the same timepoints.

**Figure 3:**
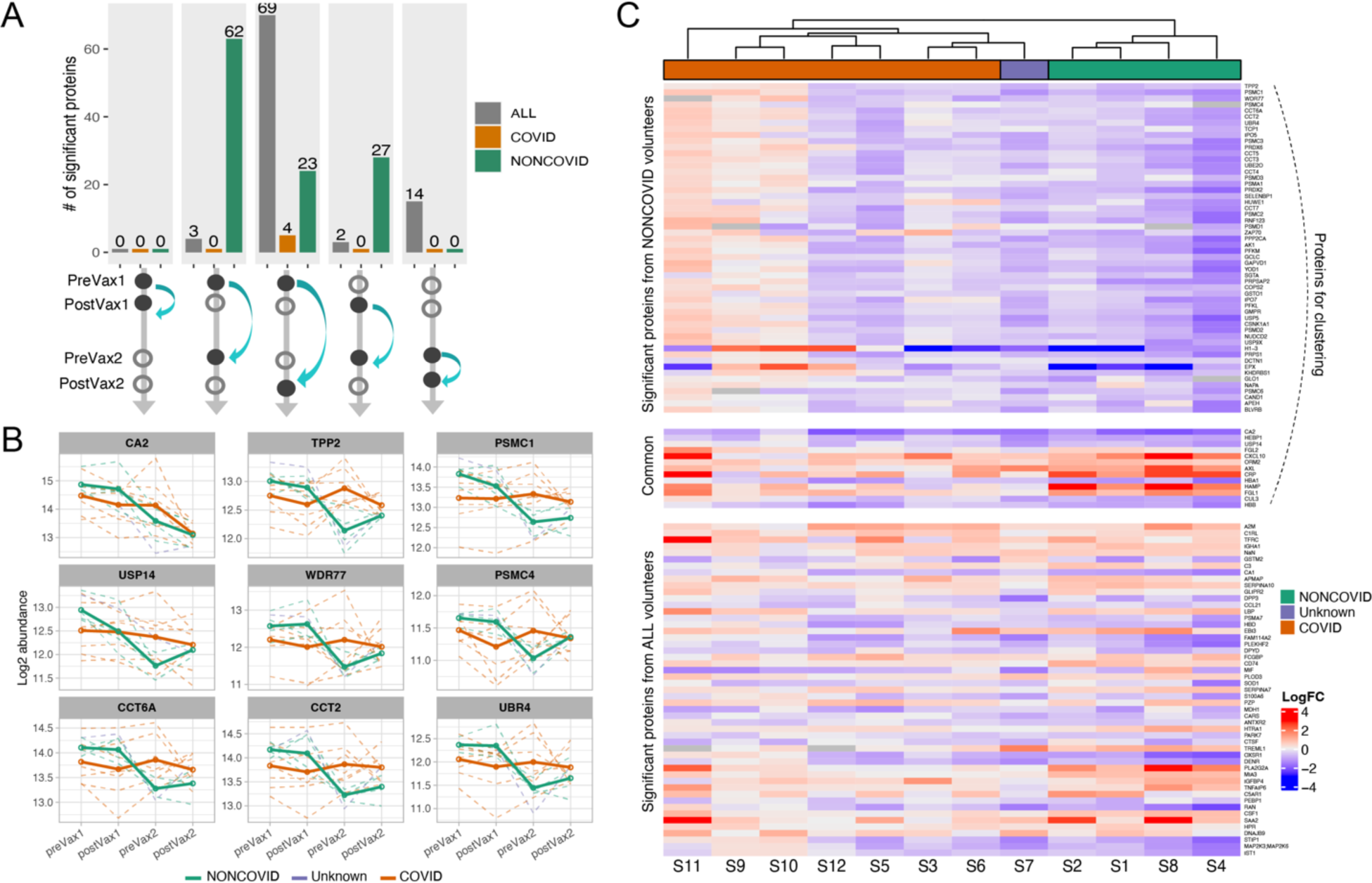
Comparison between COVID and NONCOVID individuals. (A) Number of Significant Proteins: Comparison of the number of significant proteins (adjusted p-value ≤ 0.05) using ALL 12 individuals, 7 COVID individuals, and 5 NONCOVID individuals separately. (B) The dynamics of top 9 proteins differentially regulated in NONCOVID subgroup. Dashed lines represent the log₂ protein abundance in each participant, solid lines represent the mean log₂ abundance in each subgroup: COVID (orange), NONCOVID (green), and S7 participant with cough and sore throat, not tested for COVID (purple). (C) Hierarchical Clustering based on the 77 proteins significantly regulated in NONCOVID participants. The upper heatmap block shows the proteins specific to NONCOVID, the lower block displays proteins specific in ALL individuals, and the middle section illustrates common proteins between NONCOVID and ALL. Columns represent the 12 individuals, and each cell indicates the log₂ fold change between PostVax2 and PreVax1 timepoints.

We examined whether proteins differentially regulated in the NONCOVID group exhibited different dynamics in COVID participants (**Figure 3B**). The top 9 proteins differentially regulated in the NONCOVID group had more stable dynamics in the COVID group, suggesting a weaker response to the vaccine. In contrast, the NONCOVID group showed a significant decrease in protein abundances between PostVax1 and PreVax2, followed by an increase after the second dose. Our data suggest that the mRNA vaccine can induce changes in the proteasome, as seen in the profiles of TPP2, PSMC1, PSMC4, USP14, and UBR4. Given that mRNA translates into proteins, which are then degraded by proteasome and generate epitopes loaded onto MHC class I molecules^32^, such changes are anticipated. This indicates that the vaccine prompted a significant response in NONCOVID participants, possibly establishing some initial protection even with the first dose, and the second dose enhanced the response. For instance, UBR4 exhibited decreased expression from PostVax1 to PreVax2 and increased expression at PostVax2, particularly in the NONCOVID group.

Hierarchical clustering using the abundance profiles of the 77 proteins differentially regulated in NONCOVID subgroup distinctly separated the participants from the NONCOVID and COVID groups (see **Figure 3C**). The participant S7, who exhibited symptoms, such as fever and cough, but did not undergo a COVID test, was clustered with the COVID group. The proteins significantly upregulated in the NONCOVID group were downregulated in COVID participants. However, the proteins significantly regulated only in the context of the entire cohort did not exhibit significant differences between COVID and NONCOVID groups. Proteins regulated both in NONCOVID and ALL groups had the same fold-change direction in NONCOVID and COVID but larger magnitude in the COVID group. Based on our observation, the global plasma profiles not only reveal the diversity in vaccine responses but show the potential for predicting the personal vaccination efficiency.

### Pathways enrichment in ALL and NONCOVID participants

Pathways enrichment analysis revealed 47 and 46 Gene Ontology Biological Processes significantly enriched by vaccination when comparing PostVax2 vs PreVax1 for ALL and NONCOVID participants, respectively (see **Figure 4**). 18 of these pathways were shared between ALL and NONCOVID groups and encompassed important immunity-related pathways like acute phase response, adaptive and humoral immune responses. The humoral immune response pathway was upregulated in both NONCOVID and ALL groups, indicating induction by the COVID-19 vaccine in alignment with the previous findings^6,10^.

**Figure 4:**
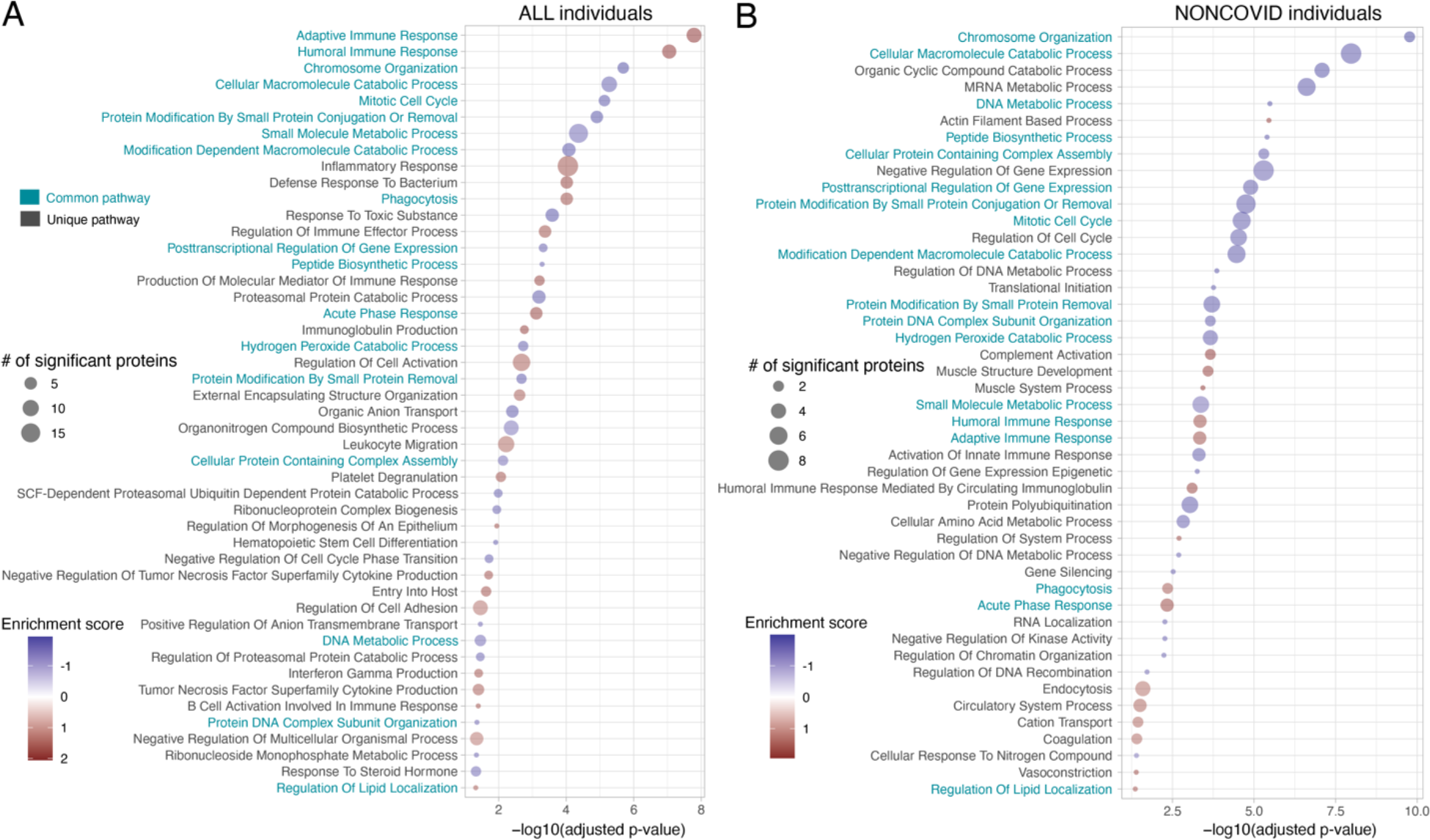
The enrichment of Gene Ontology Biological Processes in (A) ALL and (B) NONCOVID individuals. The pathways were generated based on Gene Ontology Biological Processes, categorizing all proteins with significant regulation between PostVax2 and PreVax1 samples (refer to Figure 2A for details). Pathways with a GSEA p-value of ≤ 0.05, adjusted using the Benjamini-Hochberg method, and containing more than one differentially regulated protein are listed. The pathways common between ALL and NONCOVID groups are highlighted by teal color. A positive enrichment score indicates pathway upregulation, while a negative score indicates downregulation.

In ALL group (**Figure 4A**), the upregulated proteins demonstrated significant enrichment in processes linked to inflammation and immunity. Key pathways included inflammatory response, and immunoglobulin production. Conversely, the most significantly enriched pathways identified with downregulated proteins were associated with catabolic and metabolic processes. Interestingly, the ALL group uniquely showed upregulation in the inflammatory response pathway among others. This includes cytokines (CXCL10, CSF1, SPP1, CXCL2), proteins associated with inflammation (PLA2G2A, CD44), innate immune response (HPR) and complement activation (C3), as depicted in **Supplementary Figure 6**. The activation of the inflammatory cascade critically influences the clinical manifestations of COVID-19^33^. Given its role in immune regulation and inflammation, these proteins may indirectly influence vaccine responses. The modulation of their activity could potentially impact the magnitude and quality of the immune response elicited by vaccines, affecting factors such as antigen presentation, cytokine production, and immune cell activation.

For NONCOVID individuals (**Figure 4B**), most top-ranking pathways were downregulated and associated with catabolic and metabolic processes, mirroring observations from the ALL group. Unique to the NONCOVID group were complement activation and activation of the innate immune response. Intriguingly, unlike other immunity-related pathways, activation of the innate immune response was downregulated in this group, which might be a possible mechanism of long non-coding RNA (lncRNA)-mediated suppression, a phenomenon previously reported in SARS-CoV-2 infections^34^. This potential innate immune suppression by SARS-CoV-2 mRNA vaccinations has been discussed^35^ but needs to be further investigation, especially given the novelty of this vaccine platform for infectious diseases.

While conducting differential analysis exclusively on NONCOVID individuals, we pinpointed a distinctive set of 62 proteins that exhibited differential regulation between PreVax1 and PreVax2, two months after the first dose. Notably, these proteins were found to be uniquely associated with downregulated pathways linked to the activation of innate immune response, antigen processing, and Interleukin 1 (refer to **Supplementary Table 2**). Furthermore, regulation of hydrolase activity, a mechanism believed to be significant in lung physiology and inflammation^36^, exhibited unique regulation between PreVax1 and PreVax2.

## Discussion

Several previous COVID-19 vaccination studies discussed the role of proteins, metabolites and pathways in the response to mRNA/inactivated vaccine^6,9,10^, and proposed to predict vaccine efficacy based on some of the identified biomolecules^6^. However, mRNA vaccine studies primarily focused on a limited set of inflammatory/immune proteins, while other studies employed untargeted methods to explore the inactivated vaccine. The current study, to our knowledge, is the first attempt to comprehensively explore the impact of mRNA vaccine on the plasma proteome, achieving the deepest coverage of individual proteome for COVID-19 vaccine studies.

Distinct from targeted and depletion workflows, nanoparticle-based workflow offers an enhanced breadth and depth in detection of plasma proteins. In our study, the nanoparticle-based workflow coupled with TMT labeling enables the identification of more than 3,000 proteins from only 48 human plasma samples. Within this protein set, we first recognized a group of proteins that exhibited a “dose-dependent” reaction to the vaccine, with an amplified response magnitude after the second dose. Most of these proteins exhibited similar changes in the same direction after both the first and second doses. However, the small cohort size has limited the statistical power to declare these changes after the first vaccine dose significant. These 69 proteins demonstrated a weaker response in the COVID group, suggesting a potential correlation with vaccine efficacy. Furthermore, in individuals who did not contract COVID-19, we identified another set of 77 proteins that can accurately distinguish between the COVID and NONCOVID groups. These proteins covered a diverse range of protein concentration levels, emphasizing the importance of the comprehensive plasma proteome profiling adopted in our study.

It is important to interpret the findings of this pilot study in the context of its limitations. The sample size is relatively small, comprising only five individuals in the NONCOVID group, alongside the inclusion of an Unknown volunteer, S7. Additionally, all NONCOVID individuals, except for S7, are female. Previous research has indicated that gender may influence COVID-19 outcomes, with men experiencing higher rates of hospitalization and mortality, whereas women may have a higher risk of long-term COVID-19 effects^37,38^. Evidence also suggests that the X chromosome plays a role in shaping the immune system, through its impact on proteins that may be overexpressed in women^39^. This genetic factor is thought to affect responses to viral infections and vaccinations. It is also important to acknowledge that our study was limited to plasma samples, and, while blood plasma is an accessible biofluid that could be used in diagnostics, other tissues, such as the spleen, mucosa, and lymph nodes, have to be considered to comprehensively understand the molecular mechanisms of immune responses ^40^.

Despite these limitations, our pilot study has demonstrated that nanoparticle-based unbiased plasma proteome profiling is a powerful tool to study the molecular signature of individual response to vaccination and infection. Given the accessibility of plasma samples and the scalability of the technology, follow-up studies engaging larger cohorts and incorporating samples from other tissues will provide additional insights into human response to vaccines. Advancements in MS technology and nanoparticle-based workflow will further increase the depth and sensitivity of the analysis^41^. All of these would help to illuminate the intricate dynamics of individual variability in vaccination and immunity against a spectrum of infectious diseases, including COVID-19, and highlight new avenues for improving vaccination efficiency.

## Data Availability Statement

The raw spectra data is available on ProteomeXchange under the identifier XXX. Supplementary tables include files for normalized protein abundances, results of the differential abundance analysis, and outcomes of the pathway enrichment analysis.

---

**Supplementary Table 1:** Information of the 12 individuals.

| Subject id | sex | Age | sample PreVax1 (M/D/Y) | Vax1 (M/D/Y) | sample PostVax1 (M/D/Y) | sample PreVax2 (M/D/Y) | Vax2 (M/D/Y) | sample PostVax2 (M/D/Y) | Had COVID before or during donation? | Had COVID after donation? |
| --- | --- | --- | --- | --- | --- | --- | --- | --- | --- | --- |
| S1 | F | 30 | 07/27/2021 | 07/30/2021 | 07/31/2021 | 09/27/2021 | 09/28/2021 | 09/29/2021 | Y | NONCOVID |
| S2 | F | 29 | 07/27/2021 | 08/02/2021 | 08/03/2021 | 09/24/2021 | 09/27/2021 | 09/28/2021 | N | NONCOVID |
| S3 | F | NA | 07/27/2021 | 07/30/2021 | 07/31/2021 | 09/24/2021 | 09/27/2021 | 09/28/2021 | N | COVID |
| S4 | F | NA | 08/05/2021 | 08/05/2021 | 08/06/2021 | 10/13/2021 | 10/14/2021 | 10/15/2021 | N | NONCOVID |
| S6 | F | 23 | 08/11/2021 | 08/12/2021 | 08/13/2021 | 10/14/2021 | 10/14/2021 | 10/15/2021 | N | COVID |
| S8 | F | 21 | 08/13/2021 | 08/17/2021 | 08/18/2021 | 10/13/2021 | 10/13/2021 | 10/14/2021 | N | NONCOVID |
| S9 | F | 21 | 08/17/2021 | 8/17/2021 | 08/18/2021 | 10/28/2021 | 10/28/2021 | 10/29/2021 | N | COVID |
| S10 | F | 21 | 08/17/2021 | 8/17/2021 | 08/18/2021 | 10/26/2021 | 10/26/2021 | 10/27/2021 | N | COVID |
| S5 | M | 25 | 08/10/2021 | 08/10/2021 | 08/11/2021 | 10/07/2021 | 10/07/2021 | 10/08/2021 | N | COVID |
| S7 | M | 23 | 08/12/2021 | 08/12/2021 | 08/13/2021 | 10/21/2021 | 10/21/2021 | 10/22/2021 | N | Unknown |
| S11 | M | 20 | 08/19/2021 | 08/19/2021 | 08/20/2021 | 10/19/2021 | 10/19/2021 | 10/20/2021 | Y | COVID (twice) |
| S12 | M | 23 | 08/18/2021 | 08/19/2021 | 08/20/2021 | 10/21/2021 | 10/21/2021 | 10/22/2021 | N | COVID (twice) |

**Supplementary Figure 1:**
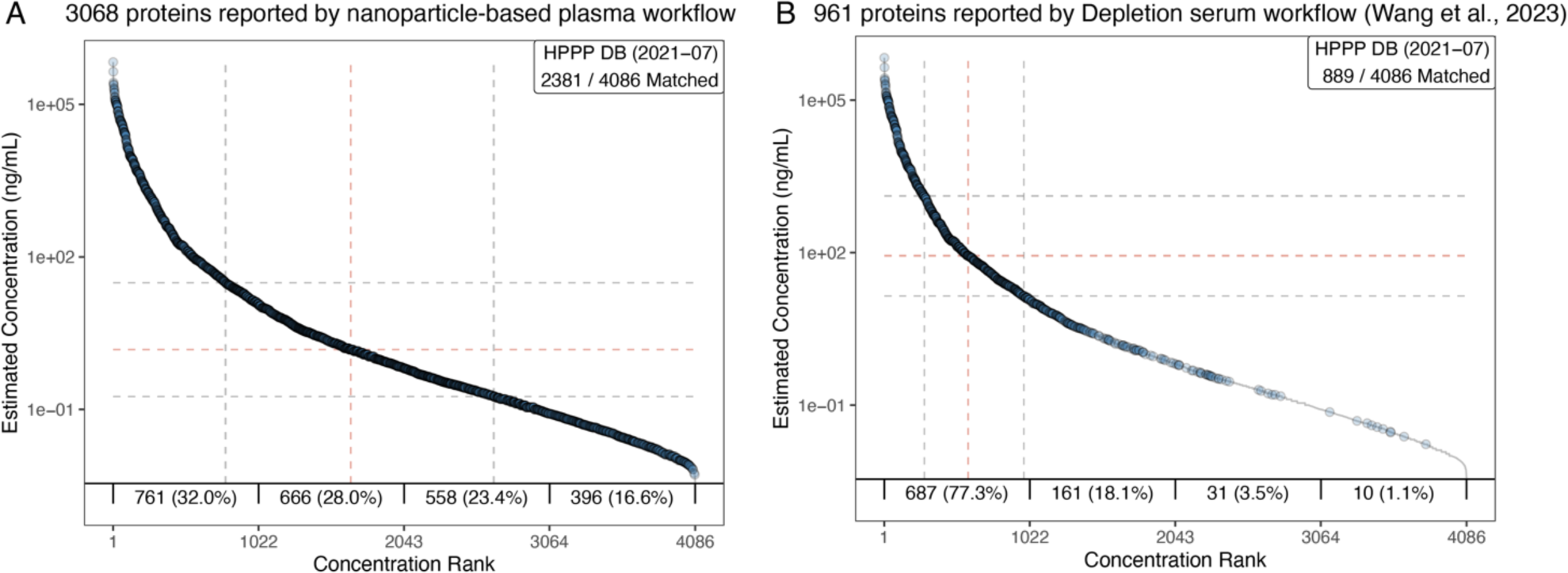
Protein overlap between our nanoparticle-based plasma workflow and an existing COVID-19 inactivated vaccine study^6^ using depletion serum workflow with proteins reported in Human Plasma Proteome Project (HPPP). We mapped the proteins identified by the two workflows to the Human Plasma Proteome Project (HPPP)^22^ protein database, respectively. Of 961 proteins reported by Depletion serum workflow^6^, 889 were mapped to the HPPP database and 4.6% were from the lower concentration range. Of 3,068 proteins reported by nanoparticle-based plasma workflow, 2,381 were mapped to the HPPP database and 40% were from the lower concentration range.

**Supplementary Figure 2:**
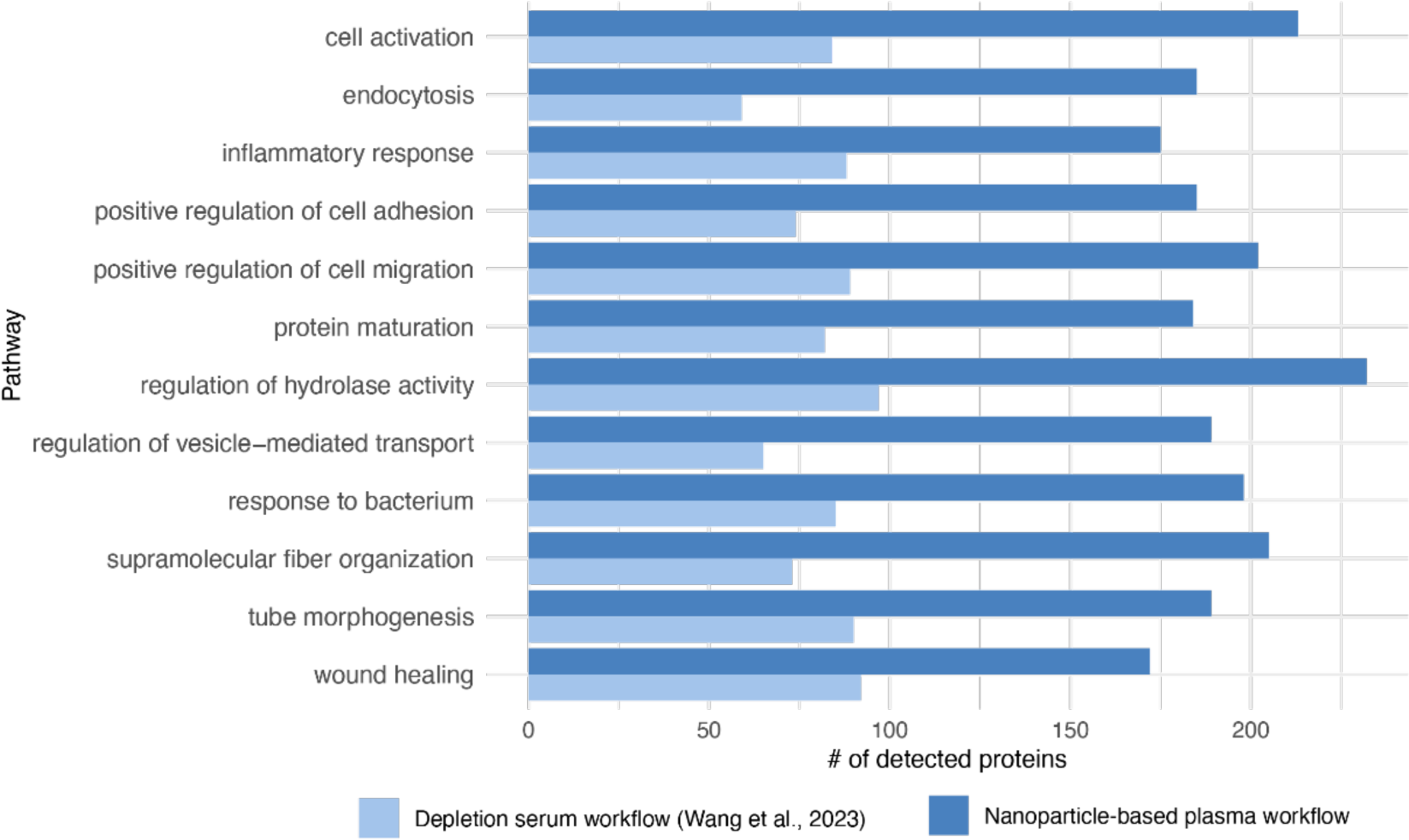
Number of proteins detected in top 12 GO biological process (GOBP) pathways. Nanoparticle-based plasma workflow detected more proteins than depletion serum workflow^6^ in each top enriched pathway.

**Supplementary Figure 3:**
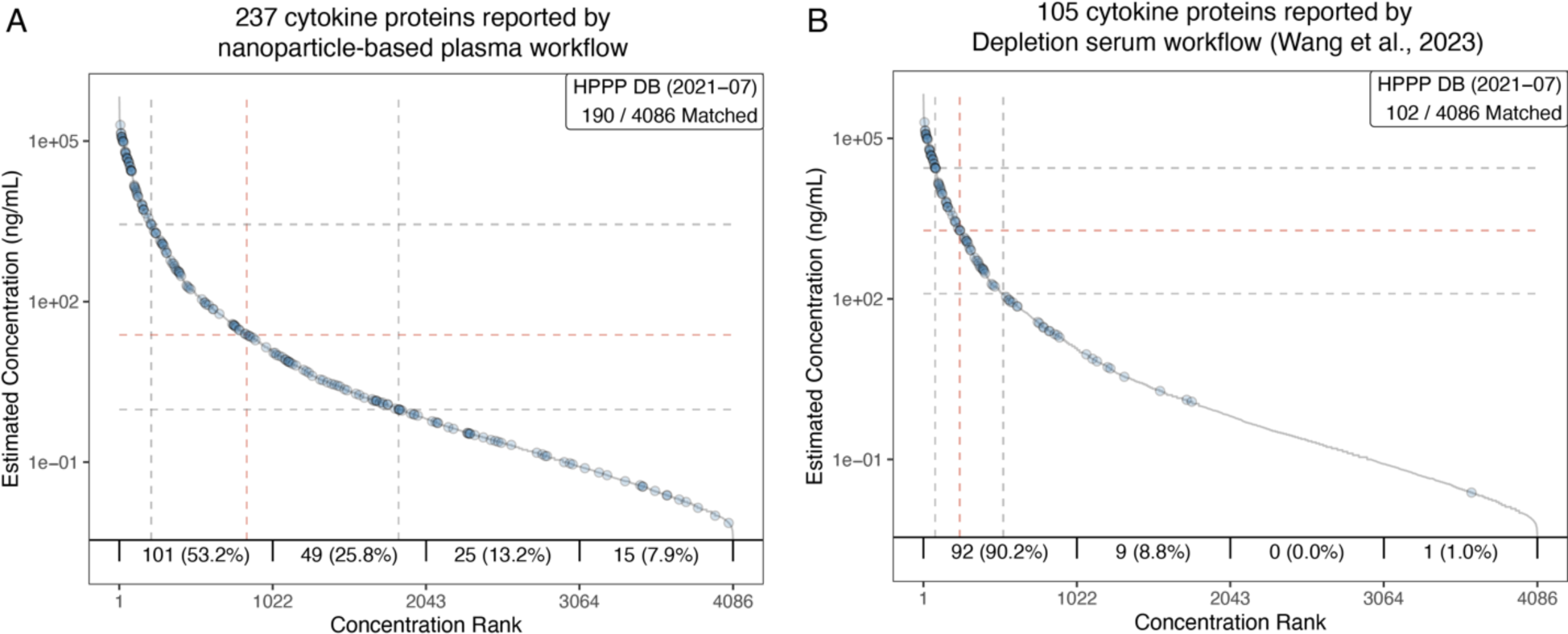
Cytokine proteins identified with nanoparticle-based plasma workflow and depletion serum workflow^6^. Of 105 cytokine proteins reported by Depletion serum workflow, 102 were mapped to the HPPP database^22^ and 1% were from the higher concentration range. Of 237 cytokine proteins reported by nanoparticle-based plasma workflow, 190 were mapped to the HPPP database and 21.1% were from the lower concentration range.

**Supplementary Figure 4:**
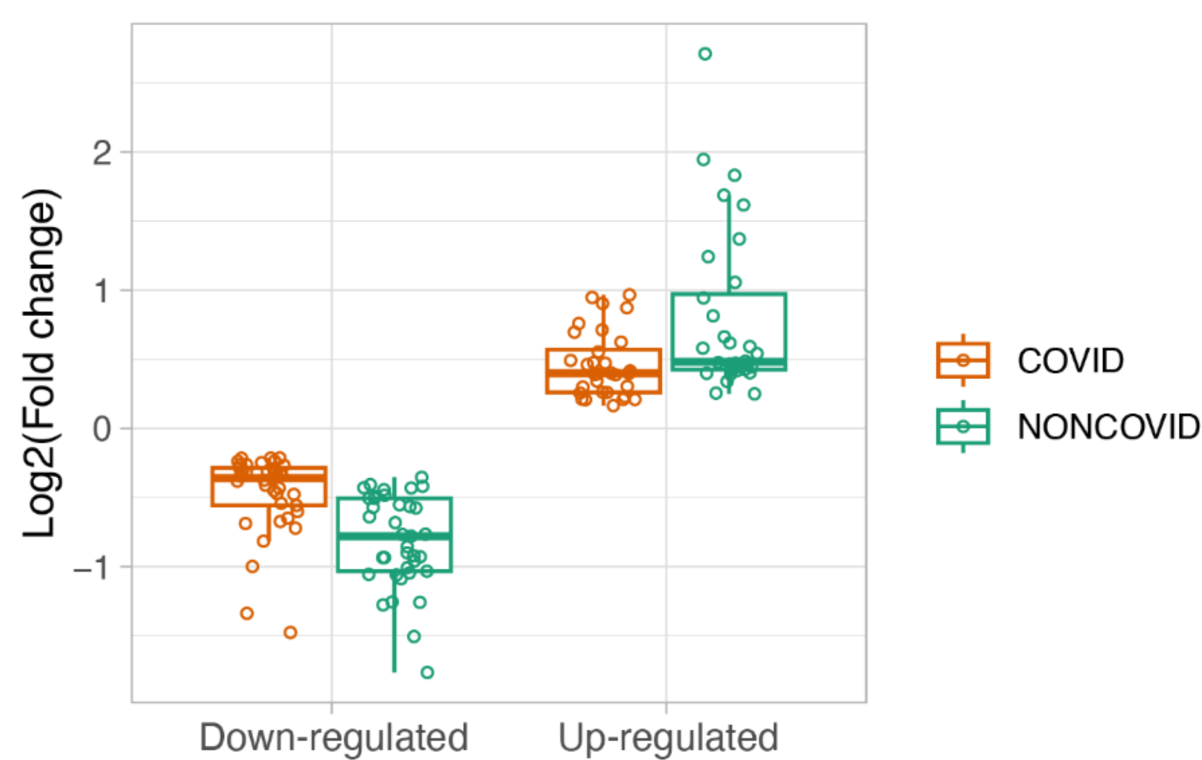
Log2 fold change of 69 significant proteins in COVID and NONCOVID subgroups. Each dot represents an individual significant protein. The 69 proteins were identified between postVax2 and preVax1 using ALL individuals, and their respective fold changes were calculated in both COVID and NONCOVID subgroups.

**Supplementary Figure 5:**
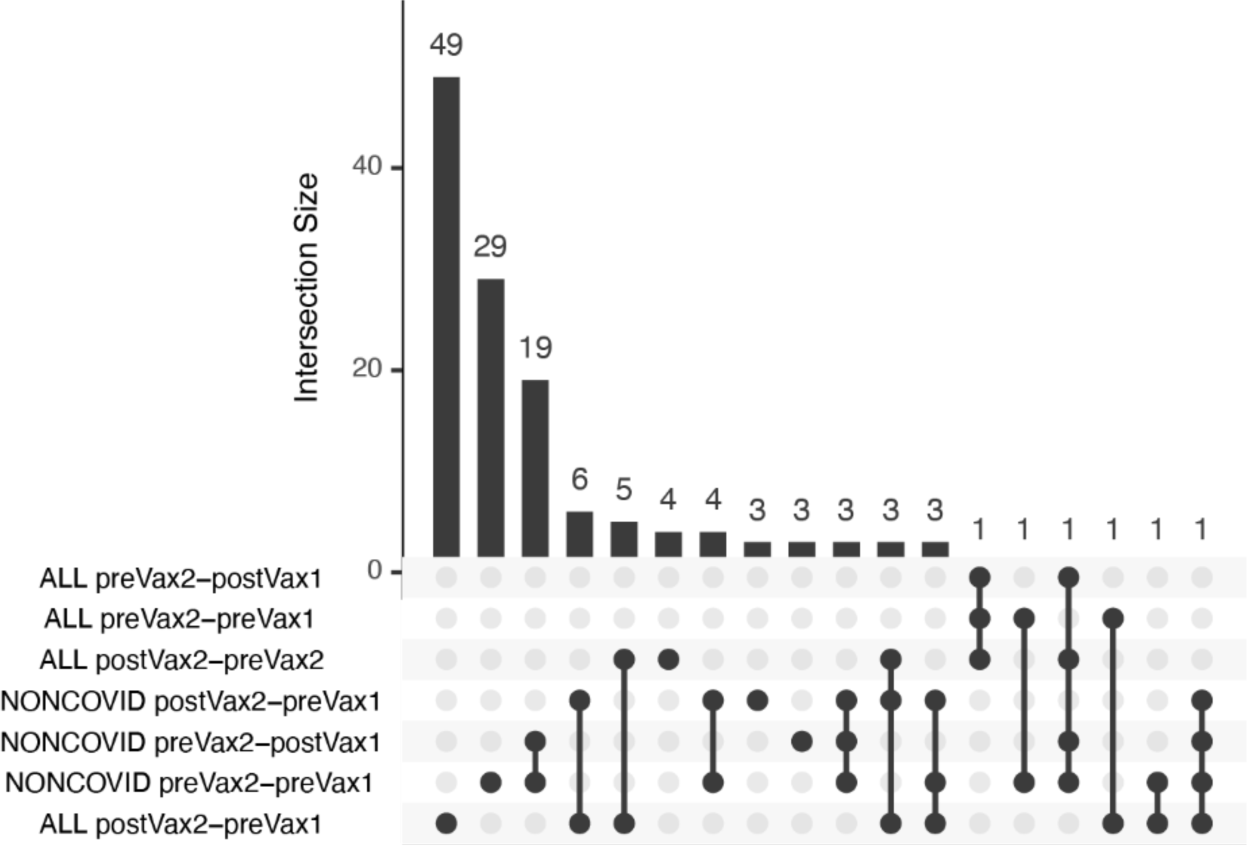
Overlap of significant proteins among different comparisons and sets of samples.

**Supplementary Figure 6:**
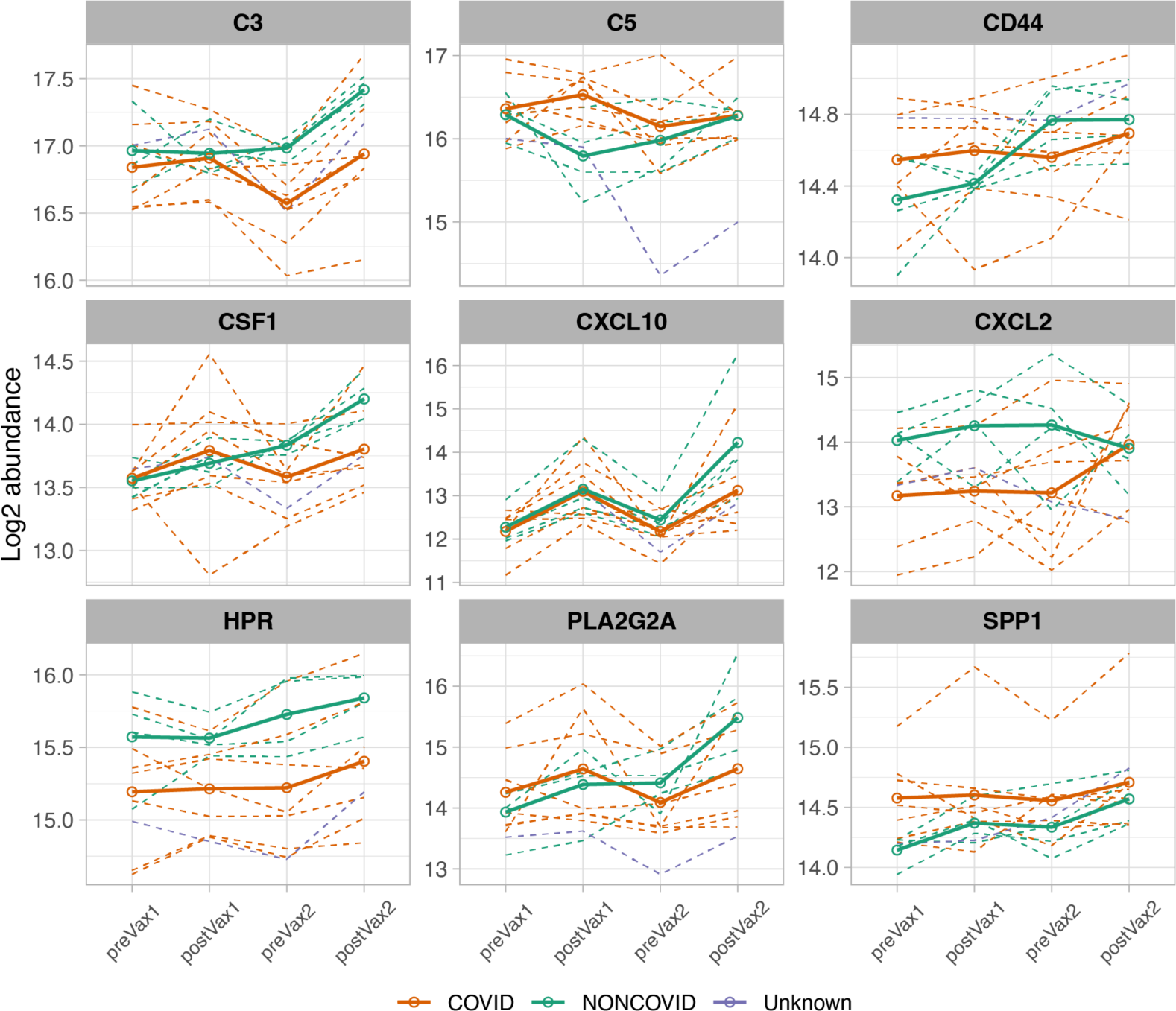
The dynamics of the 9 proteins involved in inflammatory response pathway. Dashed lines represent the log₂ protein abundance in each participant, solid lines represent the mean log₂ abundance in each subgroup: COVID (orange), NONCOVID (green), and S7 participant with cough and sore throat, not tested for COVID (purple).

**Supplementary Table 2:**
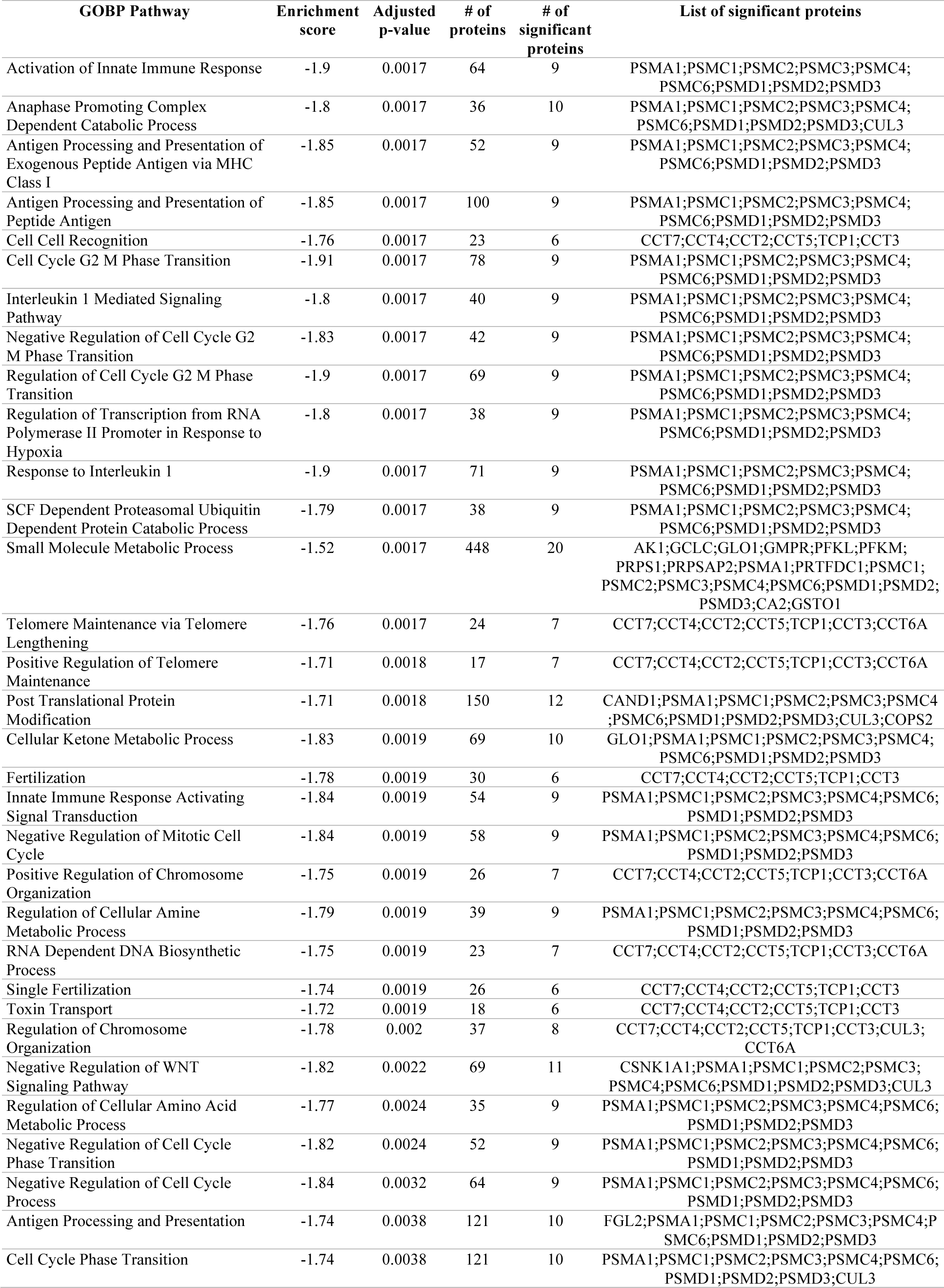
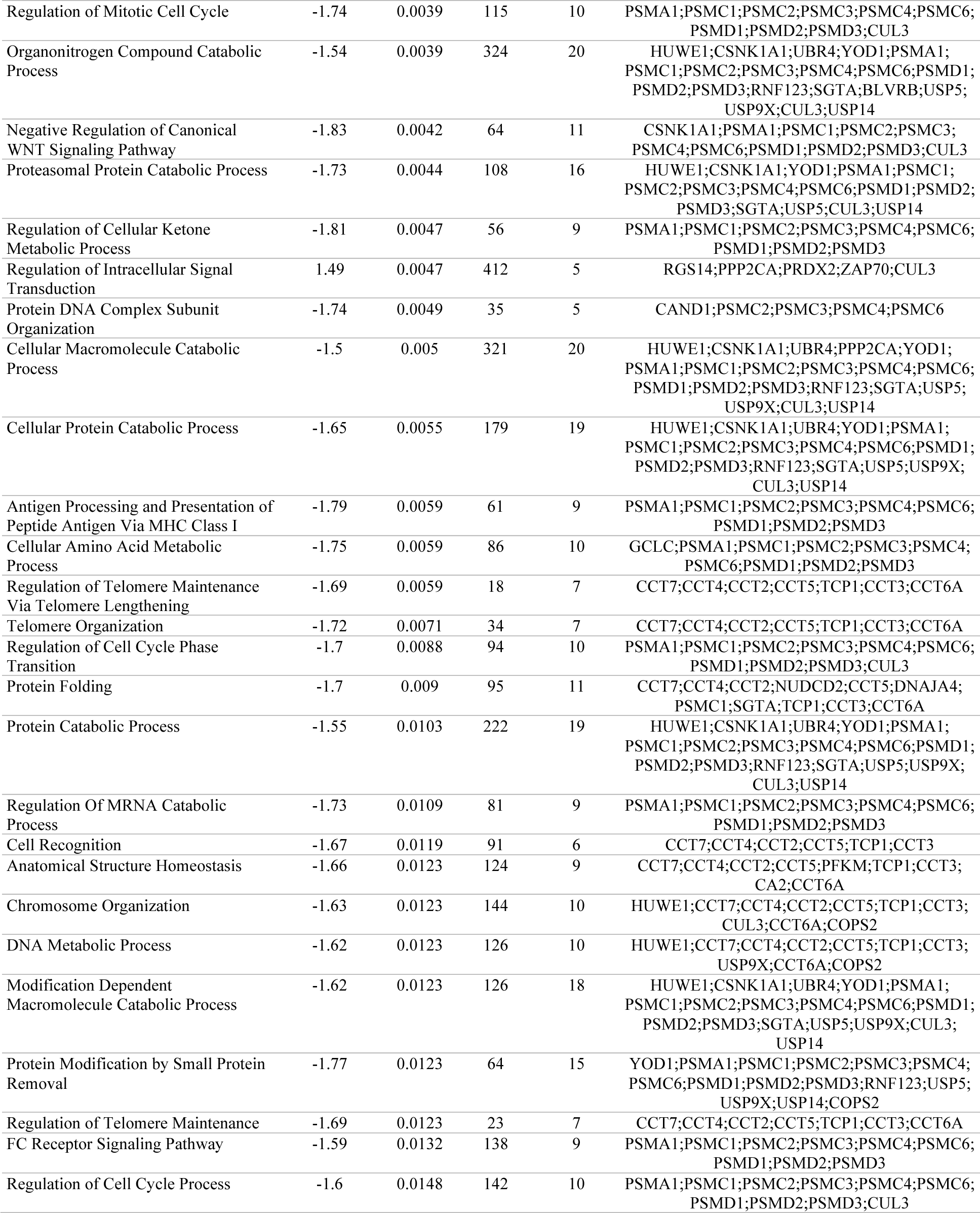
Pathway enrichment analysis based on the differentially regulated proteins in **PreVax2 and PreVax1** NONCOVID individuals. The table lists all the significantly regulated pathways with adjusted p-value <= 0.05 and regulated proteins > 5. Positive enrichment score indicates that pathways are up-regulated while negative enrichment score indicates that pathways are down-regulated. Only two out of the 58 significant pathways were upregulated.

